# Catastrophic drought effects on water quality and phytoplankton biomass in Nanwan Reservoir (Xinyang, China)

**DOI:** 10.1101/2025.04.19.649684

**Authors:** Kunjie Wu, Xin Liu, Yuan Tian, Chenxi Ju, Chaoqun Su, Xinliang Peng, Yuanye Ma, Huanan Gao, Xusheng Guo

**Affiliations:** College of Fisheries, Xinyang Agriculture and Forestry University, Xinyang, Henan 464000, China; Guangxi Key Laboratory of Aquatic Biotechnology and Modern Ecological Aquaculture, Guangxi Academy of Marine Sciences, Guangxi Academy of Sciences, Nanning, Guangxi 530007, China; College of Life and Environmental Science, Wenzhou University, Wenzhou 325035, China; Xinyang Nanwan Reservoir Affairs Center, Xinyang, Henan 464000, China; School of Environment, Tsinghua University, Beijing 100084, China

## Abstract

Climate change has led to increasingly frequent and unpredictable droughts and high-temperature events, creating extreme conditions that profoundly impact the productivity of freshwater ecosystems. This study investigates the Nanwan Reservoir, a large and deep reservoir in Xinyang, China, to evaluate the effects of extreme drought events on water quality and phytoplankton production. Field investigations were conducted during both high-temperature and drought (HTD) conditions in 2019, and during non-high-temperature and drought (NHTD) conditions in 2020. Results showed that HTD conditions significantly disrupted the thermocline and oxycline structures, leading to prolonged stratification during the HTD period. Although phosphorus concentrations remained relatively stable across both periods, nitrogen levels were markedly lower during HTD events—likely due to uptake by phytoplankton—indicating a shift in nutrient limitation from phosphorus to nitrogen. Additionally, a complex relationship between environmental variables and phytoplankton biomass was observed under HTD conditions. These findings advance our understanding of primary production responses to extreme weather events in the Nanwan Reservoir and highlight the importance of incorporating such knowledge into water resource management and ecological conservation strategies.

## Introduction

Over the past half-century, rapid global warming driven by anthropogenic activities has emerged as a major threat to the stability of aquatic ecosystems, with significant consequences for biodiversity and ecosystem functioning [1, 2]. Climate change has increased the frequency and intensity of extreme weather events, including severe droughts, typhoons, and heavy rainfall [3–5]. As such, there is an urgent need to further evaluate the impacts of human activities on aquatic ecosystems [6].

Compared to marine ecosystems, freshwater ecosystems such as lakes and reservoirs are smaller in scale but play a crucial role by providing a wide range of ecosystem services. They are also more sensitive to climate change [7]. Freshwater systems are considered both indicators and integrators of climate change, as they are closely connected to their surrounding catchment areas, and their responses to environmental changes are often immediate and pronounced [8–15]. Since surface water temperature (WT) and thermocline strength are largely regulated by air temperature, rising temperatures intensify water column stratification and prolong its duration [16, 17]. These changes in WT can significantly affect aquatic biota, leading to shifts in species composition, biomass, and productivity [18, 19].

Phytoplankton production is regulated by a range of abiotic environmental factors, including temperature, light intensity, pH, dissolved oxygen (DO), and nutrient availability. These parameters are all significantly influenced by climate change, particularly through shifts in weather patterns. Global warming alters precipitation regimes, thereby affecting nutrient loading in freshwater ecosystems [20, 21]. Moreover, rising WTs can modify the chemical characteristics of water bodies influenced by precipitation, indirectly shaping phytoplankton community composition and structure [22, 23]. These climate-induced changes complicate the underlying mechanisms driving harmful algal blooms in eutrophic lakes, often reducing the effectiveness of conventional bloom control strategies [24–26].

The combination of elevated temperatures and decreased precipitation contributes to the onset of drought conditions. However, few studies have specifically addressed the impacts of drought on lakes and reservoirs. These impacts are typically manifested as lowered water levels and reduced nutrient inflows. For instance, in Lake Taihu (China), interannual variability in cyanobacterial bloom phenology has shown a strong correlation with local climatic conditions over the past two decades [27]. Elevated temperatures have been associated with earlier onset and prolonged duration of cyanobacterial blooms [28]. Similarly, Lake Dongting (China) experienced a severe drought during the summer of 2022, during which significantly elevated phosphorus concentrations were recorded compared to historical averages [29]. This increase was likely due to phosphorus accumulation resulting from reduced water volume.

The Nanwan Reservoir is located approximately 8.5 km southwest of Xinyang City in Henan Province, China. It encompasses a watershed area of 1,100 km² and a water surface area of 75 km², with a maximum depth of 30 meters and a total storage capacity of 1.63 billion m^3^ [30, 31]. The reservoir plays a critical role in regional water supply, serving agricultural, industrial, and domestic needs, including as a primary source of drinking water for the urban population of Xinyang. Previous studies have identified reservoir bays and dam areas as hotspots for harmful algal blooms, highlighting the need for effective nutrient management strategies to mitigate bloom risks in this large, deep-water body [32]. In 2019, Xinyang experienced an exceptionally rare and prolonged extreme weather event, characterized by nearly two months of continuous high temperatures and drought from August to October. This extended period of climatic stress likely had substantial impacts on the reservoir’s ecosystem services, particularly water availability and primary productivity.

In this study, we investigated water quality parameters and phytoplankton biomass in the Nanwan Reservoir during different stages of a drought event to evaluate the effects of extreme weather on reservoir productivity. By comparing conditions during the drought period with those from a climatologically normal year, we aim to provide new insights into how such extreme events influence production dynamics in deep reservoirs. The findings contribute to the growing body of knowledge on climate-related impacts in freshwater ecosystems and offer practical guidance for the ecological management and protection of large reservoirs under changing climate conditions.

## Materials and Methods

### Investigation periods and sample collection

Between July and September from 2016 to 2021, average air temperatures in the study area ranged from 26.0 °C to 27.5 °C, with the highest and lowest averages recorded in 2019 and 2020, respectively (S1 Table). Meanwhile, total cumulative precipitation during this period was lowest in 2019 (136.5 mm) and highest in 2020 (1117.5 mm) (S1 Fig). Based on these observations, we selected two contrasting study periods for comparison: a high-temperature and drought (HTD) period from August to September 2019, and a normal period (non-high-temperature and non-drought, NHTD) from July to September 2020. Field investigations were conducted during both the early and late stages of each period. For the HTD period in 2019, field sampling was carried out on August 13 and September 30, representing the early and late stages, respectively. For the NHTD period in 2020, fieldwork was conducted on July 12 and September 27.

Three offshore sampling sites—NW1, NW2, and NW3—were established in the southern, western, and eastern regions of the Nanwan Reservoir, respectively (Fig 1). Vertical profiles of WT and DO concentrations were measured in situ using a portable DO meter (Pro 20, YSI, Ohio, USA) at intervals from 0.5 m below the surface to the bottom. Water transparency (Secchi depth, SD) was determined using a standard Secchi disk. Water samples were collected using a 5 L plexiglass water sampler at depths of 0.5 m, 5 m, 10 m, and/or 15 m, depending on seasonal water levels and site-specific depths. All collected water samples were transported to the laboratory and analyzed for water quality indicators within 24 h. Water pH was measured on-site using a digital pH meter (PHBJ-260, Lei Ci, Shanghai, China).

**Fig 1.**
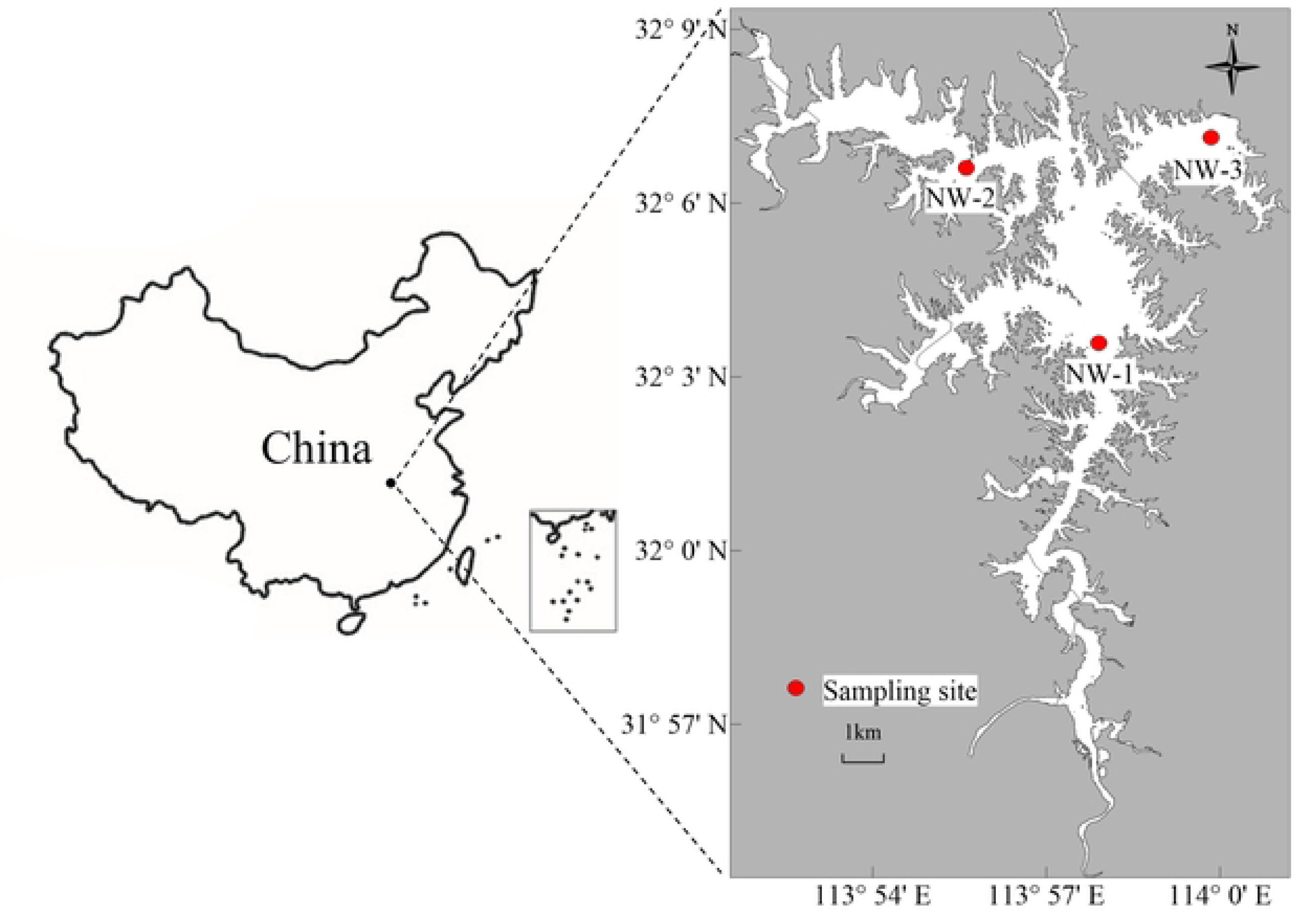
Location of Nanwan Reservoir in Xinyang, China, and three sampling sites (red circle).

### Demarcation of the thermocline and the oxycline

The thermocline forms due to a rapid decline in WT with increasing depth. It is typically defined as the water layer in which the vertical temperature gradient exceeds 1 °C per meter [33, 34]. In contrast, both the upper and lower layers adjacent to the thermocline exhibit more gradual temperature changes, with gradients of less than 1 °C per meter. In cases where the water column is shallow and no thermocline is observed, the entire water column is considered part of the upper layer of the thermocline. The oxycline, often influenced by the presence of the thermocline, is defined in this study as the water layer displaying the steepest gradient in DO concentration. In the layers above and below the oxycline, changes in DO concentration are relatively small compared to those observed within the oxycline itself [35]. Additionally, any region within the water column where DO concentrations fall below 0.2 mg/L is classified as an anoxic zone [35].

### Nutrient profiles

Nutrient concentrations and chemical oxygen demand (COD_Mn_) were determined according to the procedures described in the *Water and Wastewater Monitoring and Analysis Methods* [36]. Unfiltered water samples were used to measure total nitrogen (TN), total phosphorus (TP), and COD_Mn_. Filtered samples (0.45 μm polyester fiber membrane, 50 mm diameter; Shanghai Xingya Purification Materials Factory, Shanghai, China) were used to analyze concentrations of ammonium (NH ^+^ -N), nitrate (NO ^-^ -N), and phosphorus (PO3-4-P).

TN was extracted via potassium persulfate oxidation. Specifically, 5 mL of alkaline potassium persulfate solution was added to each sample, sealed with a ground-glass stopper, and autoclaved at 121 °C for 30 min. After cooling, 1 mL of 10% (v/v) hydrochloric acid was added. Absorbance was measured at 220 nm and 275 nm using a UV-Vis spectrophotometer (UV1700, Meixi, Shanghai, China). TP was determined by the molybdenum-antimony colorimetric method following potassium persulfate digestion. After digestion, 1 mL of 10% ascorbic acid solution was added and mixed thoroughly, followed by 2 mL of molybdate solution. The mixture was left to develop color for 15 min before measuring absorbance at 700 nm. NH ^+^ -N concentrations were determined using the Nessler’s reagent spectrophotometric method. To each sample, 1.0 mL of potassium sodium tartrate solution and 1.5 mL of Nessler’s reagent were sequentially added. After a 10-min reaction period, absorbance was measured at 420 nm. NO^-^ -N concentrations were measured via ultraviolet spectrophotometry. The ion-exchange resin was first washed, soaked in methanol overnight, and then rinsed with deionized water. To the water sample, zinc sulfate solution and sodium hydroxide were added to adjust pH to 7.0, followed by the addition of aluminum hydroxide suspension. After centrifugation, the supernatant was passed through the prepared resin adsorption column. The eluate was collected, and aliquots were transferred into cuvettes. Each aliquot received 1.0 mL of hydrochloric acid and 0.1 mL of aminosulfonic acid solution. Absorbance was recorded at 220 nm and 275 nm using a 10 mm path-length quartz cuvette. PO3-4-P in the filtrate was determined using the same molybdenum-antimony colorimetric method as TP, but without the potassium persulfate digestion step. COD_Mn_ was analyzed using the acidic permanganate method (Method A). Samples were treated with sulfuric acid and potassium permanganate solution and heated in a boiling water bath for 30 min. While still hot, a known volume of standardized sodium oxalate solution was added and thoroughly mixed. The remaining solution was immediately titrated with standardized potassium permanganate until a faint pink color persisted. The volume of potassium permanganate consumed was used to calculate the COD_Mn_ concentration. The permanganate solution was standardized prior to use with sodium oxalate.

### Eutrophication indicator

The eutrophication status of Nanwan Reservoir was assessed using the Trophic Level Index (TLI). This index employs a numerical scale ranging from 0 to 100 to categorize trophic levels. It provides a systematic framework for classifying the eutrophic status of reservoirs based on established criteria [37–39]. The TLI was calculated using the following equation [37]:

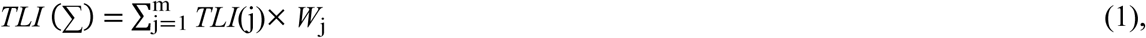

where TLI(j) denotes the nutritional state index of the *j*th substance (environmental parameters), and *W*_j_ denotes the proportion of the *j*th substance in the reservoir. In this study, TN, TP, SD, chlorophyll a (Chl. a), and COD_Mn_ were selected as the key environmental parameters. The corresponding coefficients and calculation details have been described in previous literature [37]. Under comparable nutrient conditions, a higher TLI value indicates a greater degree of nutrient enrichment, reflecting poorer water quality and more severe eutrophication (S2 Table).

### Phytoplankton biomass

Chl. a concentration is widely regarded as a reliable proxy for phytoplankton biomass in aquatic ecosystems [40–42]. In this study, Chl. a concentrations of phytoplankton assemblages were determined using the hot ethanol extraction method [43] to evaluate phytoplankton growth under different environmental conditions. For each measurement, 500 mL of water was filtered through 0.45 μm polyester fiber filters (Shanghai Xingya Purification Materials Factory, Shanghai, China).

Filters containing retained phytoplankton were stored at −20 °C until further processing. Chl. a was extracted by immersing the filters in 90% (v/v) ethanol and heated in a water bath at 80 °C for 2 min. The extracts were then kept in the dark for up to 6 h to complete pigment extraction. Absorbance was measured using a UV-Vis spectrophotometer (UV1700, Meixi, Shanghai, China). Chl. a concentration (μg/L) was calculated according to the following equation [43]:

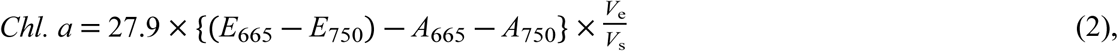

where *E*_665_ and *E*_750_ denote the absorbance values of the ethanol extract at 665 nm and 750 nm, respectively, *A*_665_ and *A*_750_ denote the absorbance values of the acidified ethanol extract at the same wavelengths, *V*_e_ (mL) is the volume of the ethanol extract, and *V*_s_ (L) is the volume of the filtered water sample.

### Statistical analysis

To assess differences between groups, Student’s t-tests were performed using the ‘t.test’ function in R. Prior to conducting t-tests, homogeneity of variances was evaluated using the ‘var.equal’ function. If variances were equal, the ‘var.equal’ argument in the ‘t.test’ function was set to TRUE; otherwise, it was set to FALSE. A Mantel test was used to evaluate the correlation between environmental variables and Chl. a concentration. In addition, stepwise regression analysis was conducted to identify key environmental factors influencing Chl. a concentrations. Given that Pearson correlation analysis indicated a strong collinearity between TP and PO3-4-P (see results), only PO3-4-P was included as the representative phosphorus variable in the stepwise regression model. All statistical analyses were performed using R software (version 4.3.0). A significance level of *P* < 0.05 was used for all tests.

## Results

### WT and DO

At all three sampling sites, surface WT at 0.5 m depth ranged from 30.6–31.1 °C during the early stage of the HTD period in 2019, and from 29.1–30.3 °C during the same stage of the NHTD period in 2020 (Fig 2a, c). A distinct thermocline was observed during both periods, with stronger stratification during the HTD period. In the early stage of HTD, the thermocline developed between 9–16 m and later stabilized at approximately 13–15 m, especially at sites NW2 and NW3 (Fig 2a). In contrast, during the early stage of NHTD, the thermocline extended more gradually from the surface down to 20 m (Fig 2c). By the late stage of both periods, surface WT declined to 24.8–26.5 °C during HTD and 25.1–25.4 °C during NHTD (Fig 2a, c). In both cases, surface temperature remained relatively constant down to ∼10 m, followed by a gradual decrease to the bottom. However, temperature declined more sharply in the HTD period (Fig 2a).

**Fig 2.**
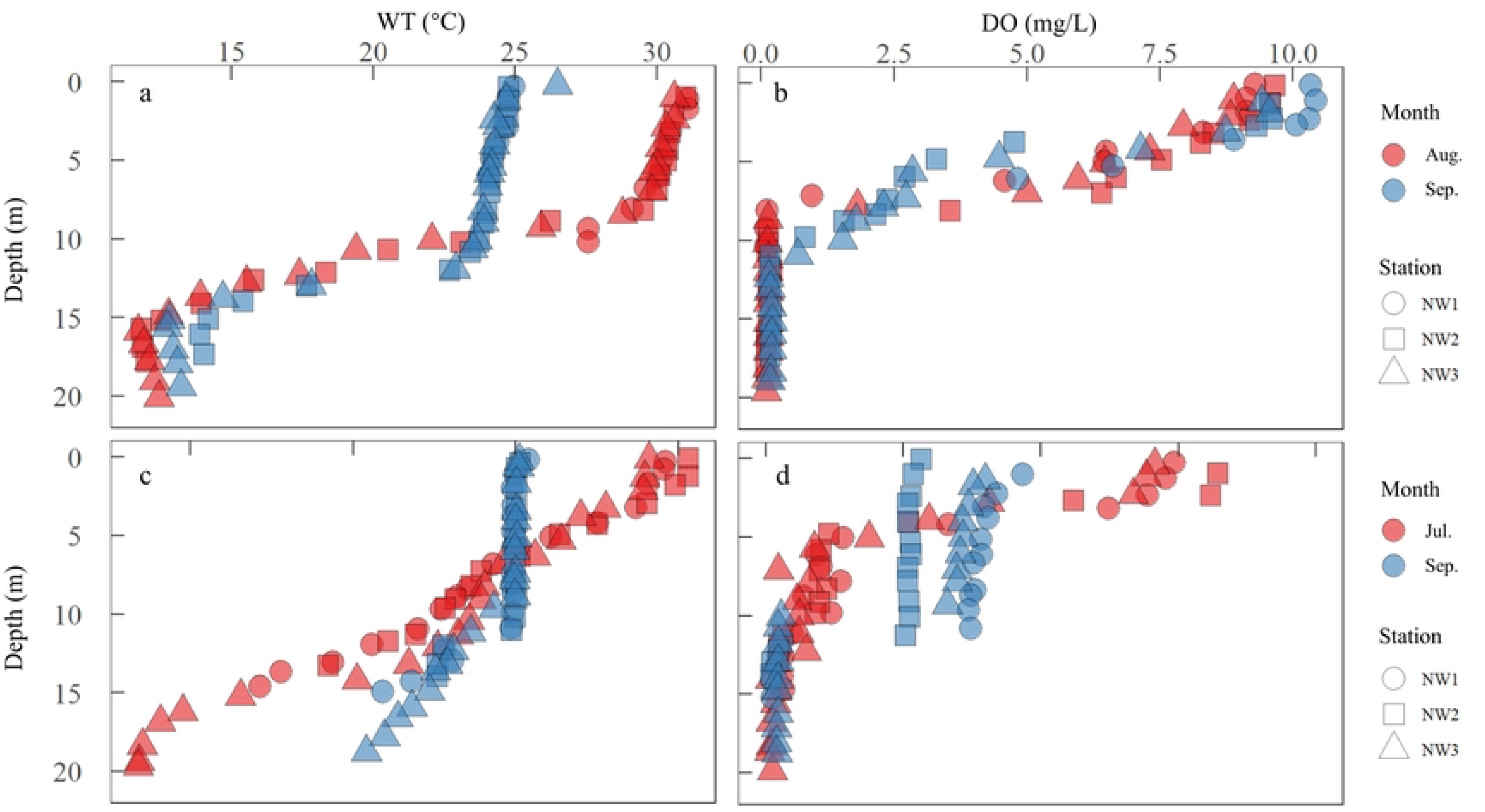
Vertical distribution of water temperature (WT, °C) and dissolved oxygen (DO) concentration (mg/L) in the Nanwan Reservoir during early (red) and late (blue) HTD period in 2019 (a, b), and early and late NHTD period in 2020 (c, d).

Vertical DO profiles during the early stages of both periods were similar, with concentrations decreasing from the surface to approximately 10 m, approaching near-zero levels at greater depths. These profiles were characterized by a clear oxycline (Fig 2b, d). In the late stage, however, differences emerged (Fig 2b, d). During late HTD, the DO profile resembled that of the early stage, with surface DO concentrations around 10 mg/L gradually declining to approximately 0.1 mg/L below 10 m (Fig 2b). In contrast, during the late NHTD period, DO concentrations ranged from 2.5– 5 mg/L between the surface and 10 m depth, then dropped sharply to <0.1 mg/L below that depth (Fig 2d).

Across all three sampling sites, the mean WT in the upper thermocline layer was 26.9 ± 3.1 °C (0–12 m) during HTD and 25.4 ± 2.3 °C (0–19 m) during NHTD (Fig 3a). In the lower thermocline layer, WT averaged 14.0 ± 3.7 °C (10–20 m) for HTD and 19.2 ± 3.9 °C (8–20 m) for NHTD (Fig 3b). For DO, concentrations in the upper oxycline layer averaged 6.88 ± 2.79 mg/L (0–10 m) during HTD and 4.24 ± 1.83 mg/L (0–12 m) during NHTD (Fig 3c). In the lower oxycline layer, DO levels were significantly lower during HTD (0.15 ± 0.03 mg/L; 8–20 m) compared to NHTD (0.56 ± 0.56 mg/L; 4–20 m) (Fig 3d). All four paired comparisons of WT and DO between HTD and NHTD periods were statistically significant (*t*-test, *P* < 0.05 for all comparisons) (Fig 3).

**Fig 3.**
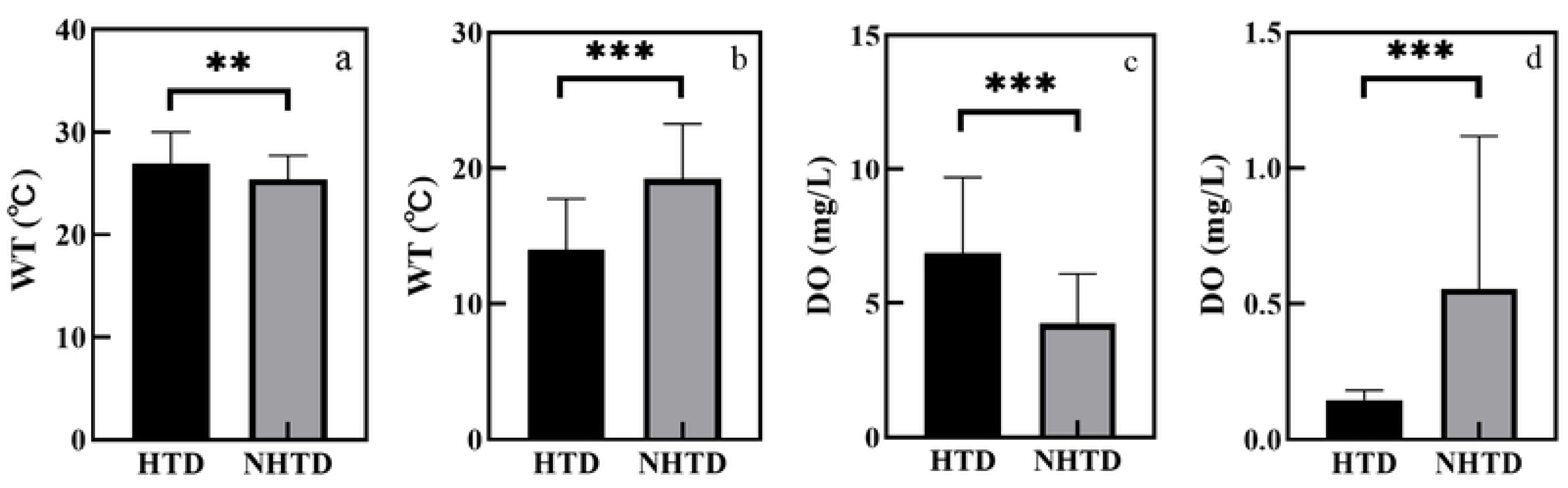
Average water temperature (WT, °C) and dissolved oxygen (DO) concentrations (mg/L) in the upper and lower layers of thermocline and oxycline during HTD and NHTD periods in Nanwan Reservoir. (a) upper layers of thermocline, (b) lower layers of thermocline, (c) upper layers of oxycline, (d) lower layers of oxycline. Results of the Student’s t-test were also provided, ** *P* < 0.01, *** *P* < 0.001.

### Water quality parameters, Chl. a and TLI indicator

Both the HTD and NHTD periods exhibited alkaline conditions, with average pH values of 7.64 ± 1.05 and 7.98 ± 0.77, respectively. A wider pH range was observed during the HTD period compared to the NHTD period (Fig 4). Water transparency, measured by SD, showed relatively narrow variation, with mean values of 1.17 ± 0.23 m for HTD and 1.29 ± 0.30 m for NHTD (Fig 4). Among nitrogen nutrients, concentrations of TN, NO-3-N and NH ^+^ -N were all significantly higher during the NHTD period than in the HTD period (*t*-test, *P* < 0.05 for all). Notably, NO-3-N concentrations were extremely low during the HTD period (0.05 ± 0.03 mg/L), while in the NHTD period, they were approximately nine times higher (0.44 ± 0.58 mg/L) (Fig 4). In contrast, phosphorus nutrients, including TP and PO3-4-P, did not differ significantly between the two periods (*t*-test, *P* > 0.05 for both). However, slightly elevated concentrations were observed during the HTD period. On average, TP concentrations were 0.04 ± 0.02 mg/L in HTD and 0.03 ± 0.02 mg/L in NHTD, while PO3-4-P concentrations were 0.03 ± 0.01 mg/L in HTD and 0.02 ± 0.01 mg/L in NHTD (Fig 4). Chl. a concentrations were slightly lower during the HTD period (9.66 ± 5.15 μg/L) compared to the NHTD period (10.42 ± 8.76 μg/L), though the difference was not statistically significant (*t*-test, *P* > 0.05). However, both COD_Mn_ and TLI values were significantly lower in the HTD period (*t*-test, *P* < 0.05). On average, COD_Mn_ was 4.00 ± 0.46 mg/L in HTD versus 7.04 ± 2.44 mg/L in NHTD, and the TLI was 43.33 ± 2.94 in HTD versus 46.57 ± 4.51 in NHTD (Fig 4).

**Fig 4.**
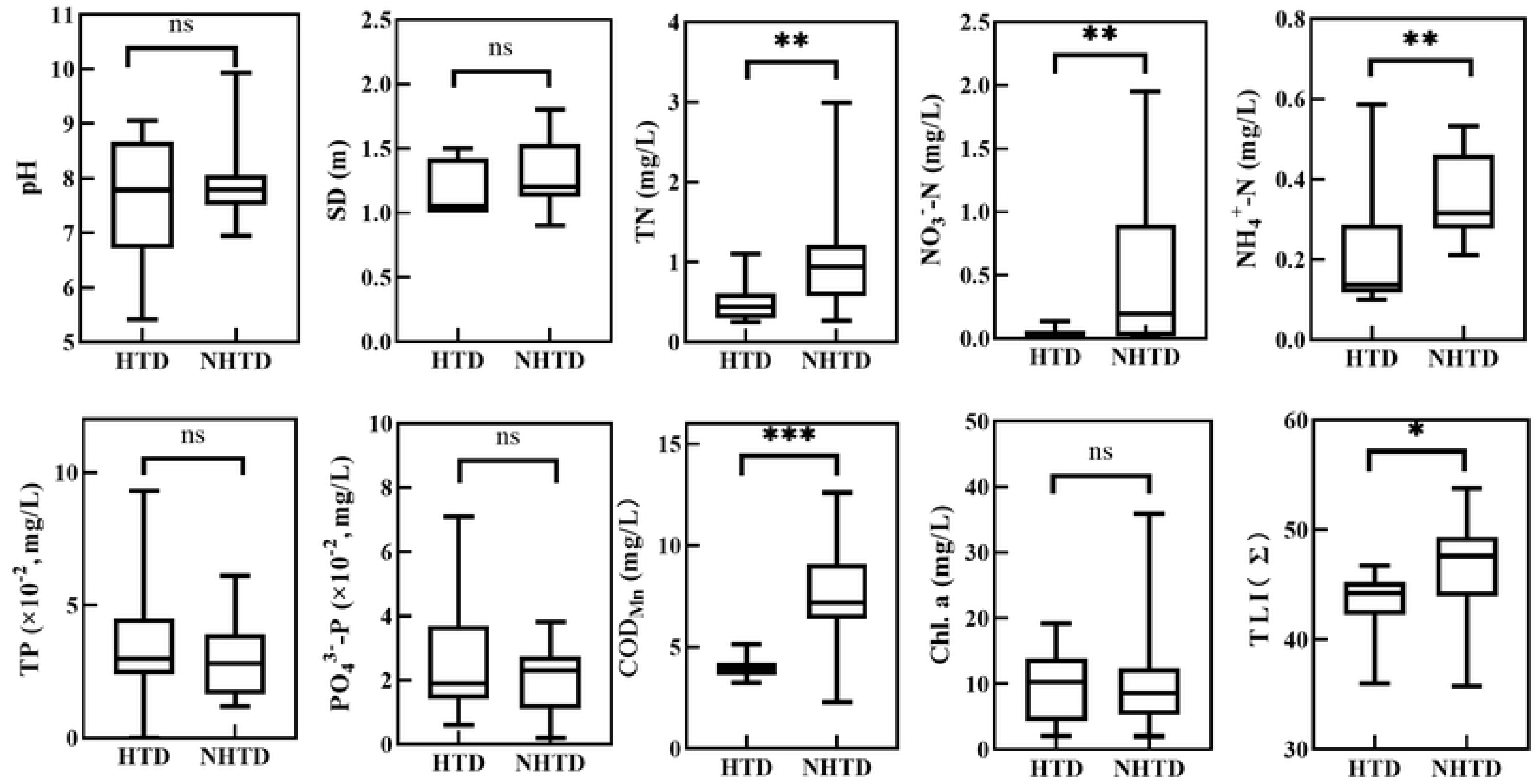
Boxplot showed pH, transparency (SD, m), nutrients (mg/L), COD_Mn_ (mg/L), Chl. a concentration (mg/L), and TLI (∑) in Nanwan Reservoir during the HTD and NHTD periods. Results of the Student’s t-test were also provided, * *P* < 0.05, ** *P* < 0.01, *** *P* < 0.001, ns no statistical difference.

### Correlation between environmental factors and phytoplankton biomass

During the HTD period, seven pairs of environmental variables exhibited statistically significant correlations (*P* < 0.05) (Fig 5a), indicating a more complex interaction network among the parameters. In contrast, only five significant correlations were observed during the NHTD period (*P* < 0.05) (Fig 5b). Specifically, during HTD, WT was positively correlated with pH (*r* = 0.75) and negatively correlated with NH ^+^ -N (*r* = -0.74) (Fig 5a). During NHTD, WT was positively correlated with both DO (*r* = 0.77) and pH (*r* = 0.70) (Fig 5b). The Mantel test revealed that Chl. a concentration was significantly correlated with several environmental factors during the HTD period, with Mantel correlation coefficients ranging from 0.22 to 0.59 (*P* < 0.05), excluding TN, NO ^-^ -N and COD_Mn_, which were not significantly correlated (Fig 5a). In contrast, during the NHTD period, Chl. a concentration showed significant correlations with only three variables—WT, DO, and pH—with stronger Mantel coefficients ranging from 0.36 to 0.72 (*P* < 0.01 for all) (Fig 5b).

**Fig 5.**
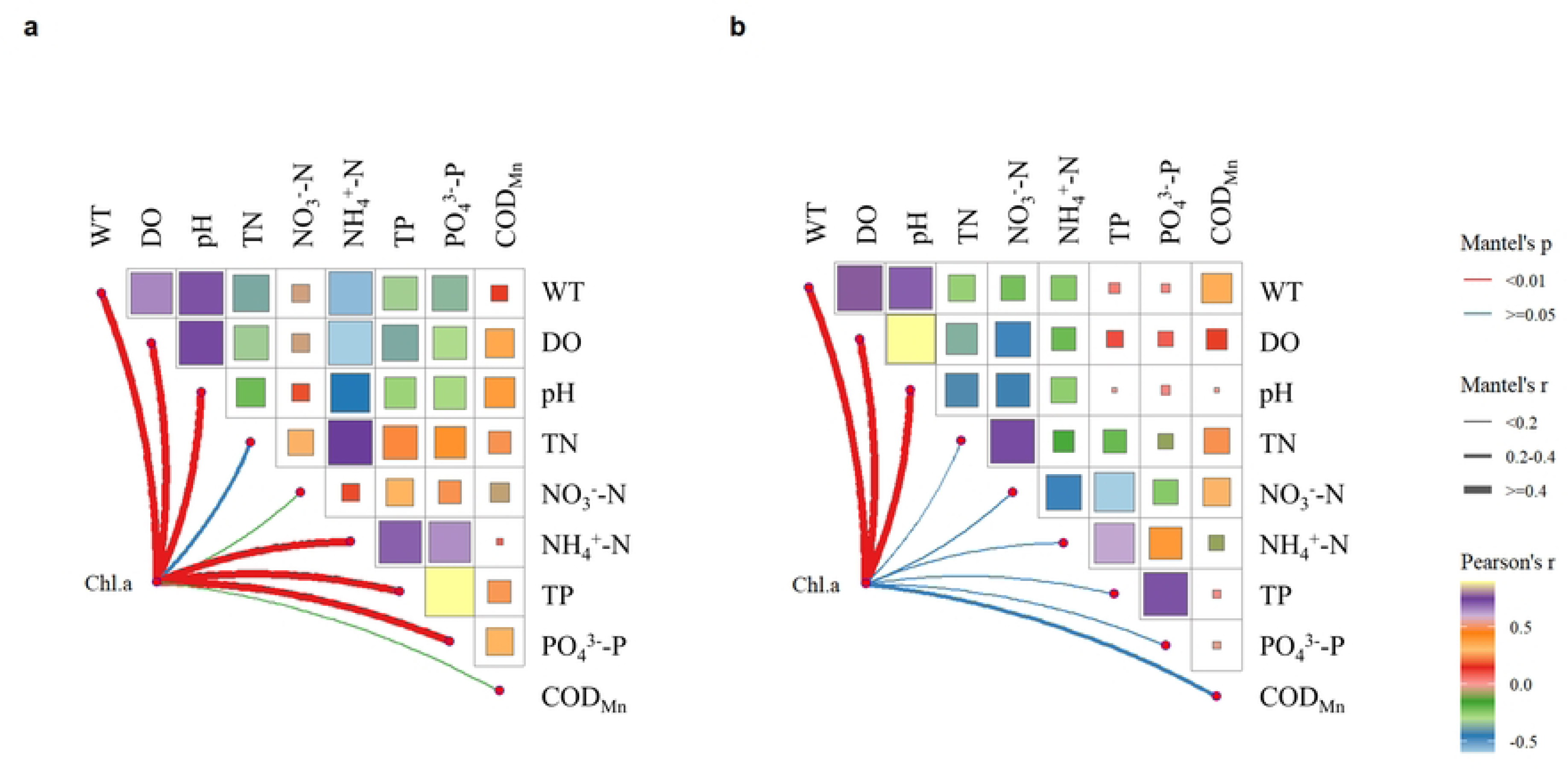
The Mantel tests showed a correlation among different environmental parameters and the relationship between Chlorophyll a (Chl. a) concentration and each tested environmental factor during HTD (a) and NHTD (b) periods in Nanwan Reservoir. * and ** indicate the statistical significance level at *P* < 0.05 and *P* < 0.01, respectively.

Stepwise regression analysis further identified the key environmental drivers of phytoplankton biomass, represented by Chl. a concentration. During the HTD period, pH and NH ^+^ -N emerged as the most influential predictors (RSS = 146.31; Akaike Information Criterion [AIC] = 43.72) (Table 1), with pH showing the strongest correlation (*P* < 0.01). In contrast, during the NHTD period, DO was the only significant predictor of Chl. a concentration (*P* < 0.01), with an RSS of 246.91 and an AIC of 55.67 (Table 1). These results suggest that the relationship between phytoplankton biomass and environmental factors shifted under the influence of catastrophic weather events, with different variables driving phytoplankton growth during drought and non-drought conditions.

**Table 1.** The best stepwise multiple regression model for the Chlorophyll a (Chl. a) concentration using environmental parameters during HTD and NHTD periods in Nanwan Reservoir. The environmental parameters included in the regression model are detailed in. **Fig. 5**.

| Period | Variables | *df* | Estimate | SE | *t* | *P* |
| --- | --- | --- | --- | --- | --- | --- |
| HTD<br>RSS = 146.31, AIC = 43.72 | Intercept | 15 | -9.14 | 7.77 | -1.78 | > 0.05 |
|  | pH |  | 2.76 | 0.90 | 3.06 | < 0.01 |
| | $\text{NH}_4^+\text{-N}$ | | -10.49 | 5.66 | -1.86 | > 0.05 |
| NHTD<br>RSS = 264.91, AIC = 55.67 | Intercept | 18 | 0.90 | 1.36 | 0.66 | > 0.05 |
|  | DO |  | 3.24 | 0.36 | 9.01 | < 0.01 |

## Discussion

A distinct thermocline is commonly observed during summer in many deep lakes and reservoirs [44, 45]. In the present study, the average temperature above the thermocline during the HTD period (26.9 °C) was 1.5 °C higher than that observed during the NHTD period (25.4 °C). This temperature increase contributed to the formation of a relatively thinner thermocline during the HTD period, likely due to enhanced thermal stratification caused by stronger density gradients in warmer surface waters [45–47]. The elevated temperatures during the HTD period led to more pronounced stratification in the Nanwan Reservoir. Given that thermal stratification creates distinct hydrodynamic conditions across water layers and inhibits vertical mixing due to temperature gradients [48], the resulting stagnation of the water column under catastrophic drought conditions may have further altered the reservoir’s physicochemical properties.

The oxycline, characterized by the steepest gradient in DO concentration, plays a crucial role in regulating oxygen availability within the water column [35]. In this study, we observed distinct patterns in the development of the oxycline between the HTD and NHTD periods in the Nanwan Reservoir. During the HTD period, the vertical DO distribution remained consistent between the early and late stages, indicating a persistently strong oxycline from August to September. In contrast, during the NHTD period, the oxycline weakened notably by September. Surface DO concentrations during the HTD period averaged 6.88 mg/L—approximately 2.64 mg/L higher than those observed during the NHTD period (4.24 mg/L). This may be partially attributed to enhanced phytoplankton photosynthetic activity under elevated temperatures [49]. However, extreme hypoxic conditions (<0.1 mg/L) were detected below 7 m during the HTD period, likely resulting from inhibited vertical mixing and increased oxygen consumption by microbial and benthic respiration [44]. Water hypoxia poses a serious threat to the ecological health of aquatic systems. For example, bottom-dwelling fish typically require DO concentrations between 4.3 and 5.7 mg/L for survival [47]. When DO levels fall below 0.7 mg/L, benthic organisms often abandon their burrows and become exposed at the sediment-water interface, resulting in elevated mortality rates [50]. In this study, DO concentrations in the bottom waters of the Nanwan Reservoir also dropped to approximately 0.1 mg/L below 10 m depth during the NHTD period, suggesting that hypoxic conditions frequently occur in deeper layers, regardless of catastrophic weather. Nevertheless, the combination of high temperatures and prolonged drought during HTD conditions appears to exacerbate bottom-water hypoxia. Sustained water column stratification and reduced mixing during such periods may compromise the ecological functioning of the reservoir, threaten aquatic productivity, and disrupt nutrient cycling, ultimately impacting biodiversity and water quality.

In the Nanwan Reservoir, concentrations of nitrogen species were significantly lower during the HTD period compared to the NHTD period. This reduction is likely attributed to decreased precipitation during the catastrophic drought, which would have reduced nutrient loading from the surrounding watershed. Similar observations have been reported in other freshwater systems; for instance, in Lake Dongting (China), TN concentrations declined by approximately 70% following the extreme drought in the summer of 2022 (0.57 mg/L) compared to the historical average from 1999–2017 (1.74 mg/L) [29]. In addition to reduced external loading, the low concentrations of inorganic nitrogen observed in surface waters during the HTD period may also reflect rapid uptake by phytoplankton stimulated by elevated temperatures. In contrast, phosphorus concentrations remained relatively stable between the HTD and NHTD periods in the Nanwan Reservoir. The Redfield molar ratio (C:P = 106 and N:P = 16) is commonly used to assess nutrient limitations for phytoplankton growth [51]. It is well established that oligotrophic and mesotrophic lakes typically exhibit phosphorus limitation [40, 52]. In this study, the average N:P molar ratio during the HTD period was 10, significantly lower than the value of 44 recorded during the NHTD period. These findings suggest a shift from phosphorus to nitrogen limitation under drought conditions in the Nanwan Reservoir. This is consistent with our previous research, which also indicated that nitrogen availability plays a key role in regulating phytoplankton production in this system [32]. Extreme drought conditions can induce nitrogen limitation even in eutrophic water bodies, particularly when anaerobic conditions develop in the water column and sediments. Phosphorus in aquatic systems is often derived from internal sources, through processes such as adsorption, flocculation, sedimentation, and release from anoxic sediments [53–56]. For example, in Lake Biwa (Japan), extreme phosphorus limitation has been observed, with N:P ratios exceeding 100 at depths of 10– 20 m during the peak phytoplankton growth season in July and October [40]. In Lake Balaton (Hungary), long-term stratification led to phosphorus release from bottom sediments, which triggered phytoplankton blooms [57]. Similarly, in Lake Taihu (China), water column stratification facilitates internal phosphorus release, thereby exacerbating bloom formation [58].

COD_Mn_ is widely recognized as an integrated indicator of the extent to which water bodies are contaminated by organic pollutants and reductive inorganic substances [36]. In the Nanwan Reservoir, a significant decrease in COD_Mn_ was observed during the HTD period, which can be attributed to reduced precipitation and, consequently, a decline in external inputs of organic and inorganic pollutants from the surrounding watershed. Similarly, the TLI was significantly lower during the HTD period compared to the NHTD period. This reduction is likely associated with decreased nutrient loading, particularly nitrogen, under drought conditions. The overall improvement in water quality observed during the HTD period may therefore be partially explained by nutrient limitation, resulting from reduced inflow of polluted runoff from the watershed. Previous studies have reported that drought conditions can promote nutrient recycling, enhance eutrophication, and degrade water quality in shallow aquatic systems [59, 60]. However, the Nanwan Reservoir, as a large and deep reservoir, exhibited a pronounced thermocline during the HTD period. This stratification likely limited the upward diffusion of internally released pollutants from the hypolimnion to the surface layers.

Nutrients can be transported from deeper layers to the surface through vertical water convection, potentially stimulating phytoplankton blooms [61, 62]. In large lakes and reservoirs, surface-layer nutrients are often rapidly depleted due to intense phytoplankton uptake [40, 52]. Consequently, the resupply of nutrients from deeper layers via water circulation plays a critical role in sustaining phytoplankton productivity [63, 64]. In the present study, phosphate limitation was not evident during the HTD period, which may explain the relatively stable phytoplankton production observed across the investigation period. Although correlation analyses revealed significant associations between phytoplankton biomass and several environmental factors, both the Mantel test and stepwise regression indicated that extreme drought conditions introduced a more complex network of relationships between environmental variables and phytoplankton growth, compared to those under normal conditions. The observed positive correlations between phytoplankton biomass and pH and/or DO concentrations in the Nanwan Reservoir likely reflect enhanced photosynthetic activity within the phytoplankton community. Overall, our findings provide evidence that catastrophic drought events can disrupt primary production processes in large, deep reservoirs. Moreover, they underscore the intricate and dynamic ecological responses of phytoplankton communities to shifting environmental conditions in freshwater ecosystems [65].

## Conclusion

HTD conditions induced significant alterations in the thermocline and oxycline structures of the Nanwan Reservoir. Our observations confirmed a prolonged period of water column stagnation during the HTD period. Although phosphorus concentrations remained relatively consistent between the HTD and NHTD periods, nitrogen concentrations dropped markedly during the HTD period—likely due to rapid uptake by phytoplankton under elevated temperatures. These results suggest that drought conditions in the Nanwan Reservoir may shift the nutrient limitation for phytoplankton growth from phosphorus to nitrogen. Additionally, we observed a more complex interplay among environmental variables and phytoplankton biomass during the HTD period. This complexity underscores the need for nonlinear dynamic models to better understand and quantify causal relationships between environmental drivers and primary production in freshwater ecosystems [66, 67].

## Acknowledgments

We thank the staff from the College of Fisheries, Xinyang Agriculture and Forestry University, for their great efforts in assisting with field sampling and supporting this study.

## Supporting information

**S1 Fig.**
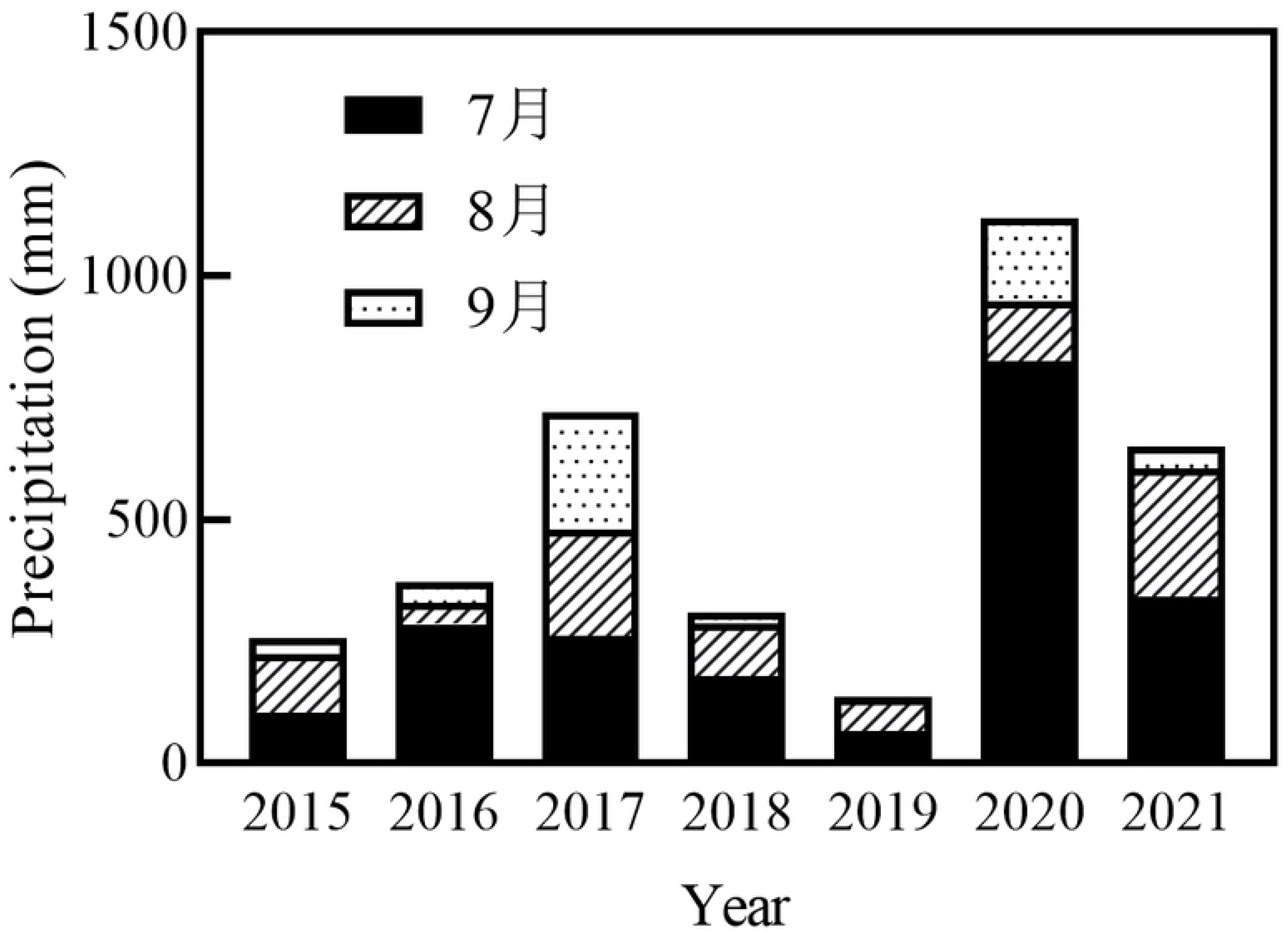
Precipitation (mm) in Xinyang City (Henan, China) during July, August, and September from 2015 to 2021. Data was provided by the Xinyang Meteorological Bureau, China.

S1 Table. The average air temperature, standard deviation (s.d.), and temperature range in Xinyang City (Henan, China) during July, August, and September from 2016 to 2021. Data was provided by the Xinyang Meteorological Bureau, China.

S2 Table. Trophic status classification of freshwater ecosystems (Jin et al. 1995; Yang et al. 2013; Huang et al. 2022).

## References

1. Dong J, Gao YN, Li GB. A review: Responses of phytoplankton communities to eutrophication and climate warming in freshwater lakers. Acta Hydrobiologica Sinica. 2016;40(03):615–623. doi: 10.7541/2016.83. (in Chinese)

2. LÜ XT, LÜ YL, Song S, Wang T. Eutrophication in cold-water lakes driven by combined effects of climate change and human activities. Acta Ecologica Sinica. 2017;37(22):7375–7386. (in Chinese)

3. Su X, Chu J, Zhang T, Jiang T, Wang G. Spatio-temporal evolution trend of groundwater drought and its dynamic response tometeorological drought in Northwest China. Water Resources Protection. 2022;38(01):34–42. doi: 10.3880/j.issn.1004-6933.2022.01.005. (in Chinese)

4. Li J, Thompson DWJ. Widespread changes in surface temperature persistence under climate change. Nature. 2021;599(7885):425–430. doi: 10.1038/s41586-021-03943-z.

5. Pörtner H-O, Roberts DC, Masson-Delmotte V, Zhai P, E. Poloczanska, Mintenbeck K, et al. Technical Summary. In: Pörtner H-O, Roberts DC, Masson-Delmotte V, Zhai P, E. Poloczanska, Mintenbeck K, et al., editors. IPCC Special Report on the Ocean and Cryosphere in a Changing Climate. Cambridge, UK and New York, NY, USA: Cambridge University Press; 2019. p. 39–69.

6. Hu B, Pan M, Shi P, Xu J, Zhang M. Effect of climate warming and eutrophication on N2O flux at water-air interface of shallow lakers. Acta Hydrobiologica Sinica. 2021;45(03):625–635. (in Chinese)

7. Chang CW, Ye H, Miki T, Deyle ER, Souissi S, Anneville O, et al. Long-term warming weakens stabilizing effects of biodiversity in aquatic ecosystems. Science advances. 2020:1–22. doi: 10.1101/2020.01.06.896746.

8. Moser KA, Baron JS, Brahney J, Oleksy IA, Saros JE, Hundey EJ, et al. Mountain lakes: Eyes on global environmental change. Global and Planetary Change. 2019;178:77–95. doi: 10.1016/j.gloplacha.2019.04.001.

9. Woolway RI, Merchant C, Hoek JVD, Azorin-Molina C, Jones ID. Northern Hemisphere atmospheric stilling accelerates lake thermal responses to a warming world. Geophysical Research Letters. 2019;46:11983–11992. doi: 10.1029/2019GL082752.

10. Woolway RI, Kraemer BM, Lenters JD, Merchant CJ, O’Reilly CM, Sharma S. Global lake responses to climate change. Nature Reviews Earth and Environment. 2020. doi: 10.1038/s43017-020-0067-5.

11. Zhu L, Zhang G, Yang R, Liu C, Yang K, Qiao B, et al. Lake variations on Tibetan Plateau of recent 40 years and future changing tendency. Bulletin of Chinese Academy of Sciences. 2019;34(11):1254–1263. doi: 10.16418/j.issn.1000-3045.2019.11.008. (in Chinese)

12. Yu X, Zhuge Y, Liu X, Du Q, Tan H. volution mechanism of dissolved oxygen stratification in a large deep reservoir. Journal of Lake Sciences. 2020;32(05):1496–1507. (in Chinese)

13. Jeppesen E, Kronvang B, Olesen JE, Audet J, Sndergaard M, Hoffmann CC, et al. Climate change effects on nitrogen loading from cultivated catchments in Europe: implications for nitrogen retention, ecological state of lakes and adaptation. Hydrobiologia. 2011;663:1–21. doi: 10.1007/s10750-010-0547-6.

14. Smol JP. The power of the past: using sediments to track the effects of multiple stressors on lake ecosystems. Freshwater Biology. 2010;55(Supplement s1):43–59. doi: 10.1111/j.1365-2427.2009.02373.x.

15. Livingstone DM. Impact of secular climate change on the thermal structure of a large temperate central European lake. Clim Change. 2003;57:205–225. doi: 10.1023/A:1022119503144.

16. Schindler DW, Bayley SE, Parker BR. The effects of climatic warming on the properties of boreal lakes and streams at the Experimental Lakes Area, northwestern Ontario. Limnology & Oceanography. 1996;41(5):1004–1017. doi: 10.4319/lo.1996.41.5.1004.

17. Wilhelm S, Adrian R. Impact of summer warming on the thermal characteristics of a polymictic lake and consequences for oxygen, nutrients and phytoplankton. Freshwater Biology. 2008;53(2):226–237. doi: 10.1111/j.1365-2427.2007.01887.x.

18. Magnuson JJ, Robertson DM, Benson BJ, Wynne RH, Livingstone DM, Arai T, et al. Historical Trends in Lake and River Ice Cover in the Northern Hemisphere. Science. 2000;289(5485):1743–1746. doi: 10.1126/science.289.5485.1743.

19. Adrian R, Walz N, Hintze T, Hoeg S, Rusche R. Effects of ice duration on plankton succession during spring in a shallow polymictic lake. Freshwater Biology. 1999;41:621–632. doi: 10.1046/j.1365-2427.1999.00411.x.

20. Bergström A-K, Jansson M. Atmospheric nitrogen deposition has caused nitrogen enrichment and eutrophication of lakes in the northern hemisphere. Global Change Biology. 2006;(12):635–643. doi: 10.1111/j.1365-2486.2006.01129.x.

21. Rogora M, Mosello R, Arisci S. The Effect of Climate Warming on the Hydrochemistry of Alpine Lakes. Water Air & Soil Pollution. 2003;148(1-4):347–361.

22. Chuai X, Chen X, Yang L, Zeng J, Miao A, Zhao H. Effects of climatic changes and anthropogenic activities on lake eutrophication in different ecoregions. International Journal of Environmental Science & Technology. 2012;9(3):503–514.

23. Sommaruga-WÖgrath S, Koinig KA, Schmidt R, Sommaruga R, Tessadri R, Psenner R. Temperature effects on the acidity of remote alpine lakes. Nature. 1997;387(6628):64–67. doi: 10.1038/387064a0.

24. Burforda MA, Careyb CC, Hamilton DP, Huisman J, Paerl HW, Wood SA, et al. Perspective: advancing the research agenda for improving understanding of cyanobacteria in a future of global change. Harmful Algae. 2020;91(21):101601. doi: 10.1016/j.hal.2019.04.004.

25. Qin B. Shallow lake limnology and control of eutrophication in Lake Taihu. Journal of Lake Sciences. 2020;32(05):1229–1243. doi: 10.18307/2020.0501. (in Chinese)

26. Ho JC, Michalak AM, Pahlevan N. Widespread global increase in intense lake phytoplankton blooms since the 1980s. Nature. 2019;574(7780):667–670. doi: 10.1038/s41586-019-1648-7.

27. Zhang M, Duan H, Shi X, Yu Y, Kong F. Contributions of meteorology to the phenology of cyanobacterial blooms: Implications for future climate change. Water research: A journal of the international water association. 2012;16:442–452.

28. Kim HG, Cha YK, Cho KH. Projected climate change impact on cyanobacterial bloom phenology in temperate rivers based on temperature dependency. Water research: A journal of the international water association. 2024;(Feb.1):249. doi: 10.1016/j.watres.2023.120928.

29. Wang D, Han J, Li R, Tang X. Nutritional characteristics in the waterbody of Lake Dongting area nutrient condition andassociated improvement measures under the extreme drought in 2022. Journal of Lake Science. 2023;35(6):1970–1978. doi: 10.18307/2023.0624. (in Chinese)

30. Yu GZ, Su H, Liu XC, Zhao CM, Gao H, Huang K. Effects of function changes of the large reservoirs in Xinyang of Henan Province on water qualities -using Nanwan Reservoir as an example. Areal research and development. 2009;28(1):115–119. in chinese

31. Fu M, Wang Z, Liang X. Floristic composition and plant community diversity of water-level fluctuation zone of Nanwan Reservoir. Desalination and water treatment. 2023;291:233–241. doi: 10.5004/dwt.2023.29438.

32. Tian Y, Ju C, Wu K, Liu X, Zhang H, Guan J, et al. Morphological characteristics and annual population dynamics of Microcystis (Cyanobacteria) in Nanwan Reservoir (Xinyang, China). Water. 2024;16:3569. doi: 10.3390/w16243569.

33. Liu X, Huang T, Li N, Yang S, Li Y, Xu J, et al. Algal bloom and mechanism of hypoxia in the metalimnion of the Lijiahe Reservoir during thermal stratification. Environmental Science 2019;40(05):2258–2264. doi: 10.13227/j.hjkx.201809026. (in Chinese)

34. Zhang Y, Chen W, Yang D, Huang W, Jiang J. Monitoring and analysis of thermodynamics in Tianmuhu Lake. Advances in water science. 2004;15(01):61–67. doi: 10.14042/j.cnki.32.1309.2004.01.012. (in Chinese)

35. Qiu XP, Huang TL, Zeng MZ. Responses of dissolved oxygen on thermal stratification and eutrophication in lakes and reservoirs—An example in Zhoucun Reservoir in Zaozhuang City. China Environmental Science. 2016;36(05):1547–1553. (in Chinese)

36. Wei F. Methods for monitoring and analyzing water and wastewater, 4th ed. Beijing, China: China Environmental Science Press; 2002.

37. Huang W, Liang N, Zhou L, Lu J. Study on the water eutrophication evolution characteristics of Junshan Lake. Water Supply. 2022;22(12):8698–8707.

38. Yang ML, Hu ZJ, Liu QG, Ren LP, Chen LS, Li PP. Evaluation of water quality by two trophic state indices in Lake Qiandaohu during 2007-2011. Journal of Shanghai Ocean University. 2013;22(2):240–245. (in Chinese)

39. Jin X, Liu S, Zhang Z. Chinese Lake Environment. Bei Jing: China Ocean Press; 1995. 278 p. (in Chinese)

40. Liu X, Yi R, Ohashi M, Maruo M, Iseri Y, Hao A, et al. Spatio-temporal variations of orthophosphate in a large mesotrophic lake and implications for phytoplankton productivity. Inland waters. 2025. doi: 10.1080/20442041.2025.2449763.

41. Liu X, Dur G, Ban S, Sakai Y, Ohmae S, Morita T. Planktivorous fish predation masks anthropogenic disturbances on decadal trends in zooplankton biomass and body size structure in Lake Biwa, Japan. limnology and oceanography. 2020;65:667–682. doi: 10.1002/lno.11336.

42. McQuatters-Gollop A, Raitsos DE, Edwards M, Pradhan Y, Mee LD, Lavender SJ, et al. A long-term chlorophyll data set reveals regime shift in North Sea phytoplankton biomass unconnected to nutrient trends. Limnology and oceanography. 2007;52(2):635–648. doi: 10.4319/lo.2007.52.2.0635.

43. Chen Y, Gao X. Comparison of two methods for Phytoplankton Chlorophyll-a Concentration Measurement. Journal of Lake Science. 2000;12(2):185–188. (in Chinese)

44. Wu Z, Liu M, Lan J, He J, Yu Z. Vertical distribution of phytoplankton and physic-chemical characteristics in the lacustrine zone of Xin’anjiang Reservoir (Lake Qiandao) in subtropic China during summer stratification. Journal of Lake Sciences. 2012;24(03):460–465. doi: 10.3969/j.issn.1003-5427.2012.03.019. (in Chinese)

45. Lin J, Su Y, Zhong H, Chen Y, Li Y, Lin H. Vertical distribution of phytoplankton in a eutrophic reservoir, Shanzi Reservoir (Fujian) during summer stratification. Journal of Lake Sciences. 2010;22(02):244–250. doi: 10.18307/2010.0214. (in Chinese)

46. Li X. Limnology. Beijing: Science press; 2013. (in Chinese)

47. Diaz RJ, Rosenberg R. Spreading dead zones and consequences for marine ecosystems. Science. 2008;321(5891):926–929. doi: 10.1126/science.1156401.

48. Komatsu E, Fukushima T, Harasawa H. A modeling approach to forecast the effect of long-term climate change on lake water quality. Ecological Modelling. 2007;209:351–366. doi: 10.1016/j.ecolmodel.2007.07.021.

49. Raven JA, Geider RJ. Temperature and algal growth. New phytologist. 1988;110(4):441–461. doi: 10.1111/j.1469-8137.1988.tb00282.x.

50. Diaz R, Rosenberg R. Marine benthic hypoxia: A review of its ecological effects and the behavioural response of benthic macrofauna. Oceanography and marine biology 1995;33:245–303.

51. Redfield AC. The biological control of chemical factors in the environment. American scientist. 1958;46(3):230A–221.

52. Wetzel RG. Limnology: lake and river ecosystems: gulf professional publishing; 2001.

53. Duhamel S, Nogaro G, Steinman Alan D. Effects of water level fluctuation and sediment–water nutrient exchange on phosphorus biogeochemistry in two coastal wetlands. Aquatic Sciences. 2016.

54. Schindler DW, Carpenter SR, Chapra SC, Hecky RE, Orihel DM. Reducing Phosphorus to Curb Lake Eutrophication is a Success. Environmental Science & Technology. 2016;50(17):8923. doi: 10.1021/acs.est.6b02204.

55. Chen J, Xu H, Zhan X, Zhu G, Qin B, Zhang Y. Mechanisms and research methods of phosphorus migration and transformation acrosssediment-water interface. Journal of Lake Science. 2019;31(04):907–918. doi: 10. 18307 /2019. 0416. (in Chinese)

56. Wetzel R. Limnology: lake and river ecosystems. Houston (TX): Gulf Professional Publishing; 2001.

57. Istvánovics V, Honti M, Torma P, Kousal J. Record-setting algal bloom in polymictic Lake Balaton (Hungary): A synergistic impact of climate change and (mis)management. Freshwater Biology. 2022;67:1091–1106. doi: 10.1111/fwb.13903.

58. Qin B, Deng J, Shi K, Wang J, Brookes J, Zhou J, et al. Extreme climate anomalies enhancing Cyanobacterial blooms in Eutrophic Lake Taihu, China. Water Resources Research. 2021;57(7):1–12. doi: 10.1029/2020WR029371.

59. Braga GG, Becker V, Rodrigues de Mendonça J, Pinheiro de Oliveira JN, de Medeiros Bezerra AF, Torres LM, et al. Influence of extended drought on water quality in tropical reservoirs in a semiarid region. Acta Limnologica Brasiliensia. 2015;27(1):15–23. doi: 10.1590/S2179-975X2214.

60. Beklioglu M, Romo S, Kagalou I, Quintana X, Bécares E. State of the art in the functioning of shallow Mediterranean lakes: workshop conclusions. Hydrobiologia. 2007;584(1):317–326. doi: 10.1007/s10750-007-0577-x.

61. Qin B, Paerl HW, Brookes JD, Liu J, Jeppesen E, Zhu G, et al. Why Lake Taihu continues to be plagued with cyanobacterial blooms through 10 years (2007–2017) efforts. Science Bulletin. 2019;64(6):354–356. doi: 10.1016/j.scib.2019.02.008.

62. Chang T, Li J, Chen W, Guo X, Liu W, Li Y, et al. Research progress in theories and approaches for lake ecosystem restoration. Advances in Water Science. 2024:1–15. (in Chinese)

63. Smith VH, Tilman GD, Nekola JC. Eutrophication: impacts of excess nutrient inputs on freshwater, marine, and terrestrial ecosystems. Environmental Pollution. 1999;100(1-3):p.179–196. doi: 10.1016/S0269-7491(99)00091-3.

64. Xie P, Chen J, Liu J. A regime shift from cyanobacterial steady state to non-cyanobacterial one by using non-traditional biomanipulation-A whole lake testing experiment in Lake Donghu, Wuhan. Journal of Lake Science. 2023;35(1):1–11. doi: 10.18307/2023.0199. (in Chinese)

65. Li P, Shi W, Liu Q, Yu Y, He G, Chen L, et al. Spatial and temporal distribution patterns of chlorophyll-a and the correlation analysis with environmental factors in Lake Qiandao. Journal of Lake Sciences. 2011;23(4):568–574. doi: 10.18307/2011.0412. (in chinese)

66. Sugihara G, May R, Ye H, Hsieh C-h, Deyle E, Fogarty M, et al. Detecting causality in complex ecosystems. science. 2012;338(6106):496–500.

67. Liu X, Dur G, Ban S, Sakai Y, Ohmae S, Morita T. Planktivorous fish predation masks anthropogenic disturbances on decadal trends in zooplankton biomass and body size structure in Lake Biwa, Japan. Limnology and Oceanography. 2020;65(3):667–682.

